# High-confidence placement of difficult-to-fit fragments into electron density by using anomalous signals - a case study using hits targeting SARS-CoV-2 non-structural protein 1

**DOI:** 10.1101/2023.06.16.545251

**Authors:** Shumeng Ma, Vitaliy Mykhaylyk, Matthew W. Bowler, Nikos Pinotsis, Frank Kozielski

**Affiliations:** School of Pharmacy, University College London, London, 29-39 Brunswick Square, London WC1N 1AX, United Kingdom; Diamond Light Source, Harwell Campus, Didcot OX46PA, UK; European Molecular Biology Laboratory, Grenoble, France; Institute of Structural and Molecular Biology, Birkbeck College, WC1E 7HX London, United Kingdom

**Keywords:** SARS-CoV-2, Covid-19, non-structural proteins, nsp1, tuneable wavelength, fragment orientation

## Abstract

The identification of multiple simultaneous orientations of small molecule inhibitors binding to a protein target is a common challenge. It has recently been reported that the conformational heterogeneity of ligands is widely underreported in the Protein Data Bank, which is likely to impede optimal exploitation to improve affinity of these ligands^1^. Significantly less is even known about multiple binding orientations for fragments (< 300 Da) although this information would be essential for subsequent fragment optimisation using growing, linking or merging and rational structure-based design. Here we use recently reported fragment hits for the SARS-CoV-2 non-structural protein 1 (nsp1) N-terminal domain to propose a general procedure for unambiguously identifying binding orientations of 2-dimensional fragments containing either sulphur or chloro substituents within the wavelength range of most tunable beamlines. By measuring datasets at two energies, using a tuneable beamline operating in vacuum and optimised for data collection at very low X-ray energies, we show that the anomalous signal can be used to identify multiple orientations in small fragments containing sulphur and/or chloro substituents or to verify recently reported conformations. Although in this specific case we identified the positions of sulphur and chlorine in fragments bound to their protein target, we are confident that this work can be further expanded to additional atoms or ions which often occur in fragments. Finally, our improvements in the understanding of binding orientations will also serve to advance the rational optimisation of SARS-CoV-2 nsp1 targeting fragment hits.

## Introduction

Although various biophysical methods exist, X-ray crystallography is considered the gold standard in fragment-based drug discovery (FBDD) as it can instantaneously provide detailed molecular information about the exact binding configuration of fragments^1, 2^. This can then be exploited for subsequent rational structure-based drug design^3^. However, fragments are small (< 300 Da) and due to their size, it can be challenging to fit these into experimental electron density, a task that can be particularly aggravated at low resolution. However, even at high resolution (< 2.0 Å), fragments can still be ambiguous to fit, in particular when they are mostly planar and substituents have not yet been optimised for stronger binding and defined interactions with residues in the binding site. Electron density from fragment binding to proteins can also be incomplete, or weak due to low occupancy of the fragment hit, further complicating interpretation of the experimental map. In the best cases there may only exist a single binding conformation but several orientations may be present at the same time^4, 5^. However, before engaging in structure-based rational design of fragment hits, it is essential to possess precise and accurate knowledge about the exact binding conformation to apply rational drug design techniques.

Recently, we reported our first fragments hits targeting SARS-CoV-2 non-structural protein 1 (nsp1) using fragment-based screening via X-ray crystallography^6, 7^. Nsp1 is a two domain protein with a short C-terminal domain, which is involved in the shutdown of host expression by blocking the entry of host mRNA in the ribosome^8^. The C-terminal domain is connected via a linker region with the larger N-terminal domain, which is thought to be involved in circumventing the shutdown of expression of viral mRNA^9^ and stabilising the binding of the C-terminus to the ribosome^10^. Therefore, nsp1 is considered as a potential target for antiviral drug discovery.

All fragment hits were essentially planar (Fig 1A) with non-optimised substituents allowing, in principle, up to four distinct binding orientations in the electron density (Fig 1B). Although working at resolutions between 1.1 and 1.4 Å, calculation of R_free_ values, taking into account atomic B-factors of fragments and careful inspection of electron density for all four configurations did not lead to an absolutely confident identification of a single correct orientation of the fragment hits. Therefore, the most reasonable fragment conformations have been reported^6, 7^.

**Fig 1.**
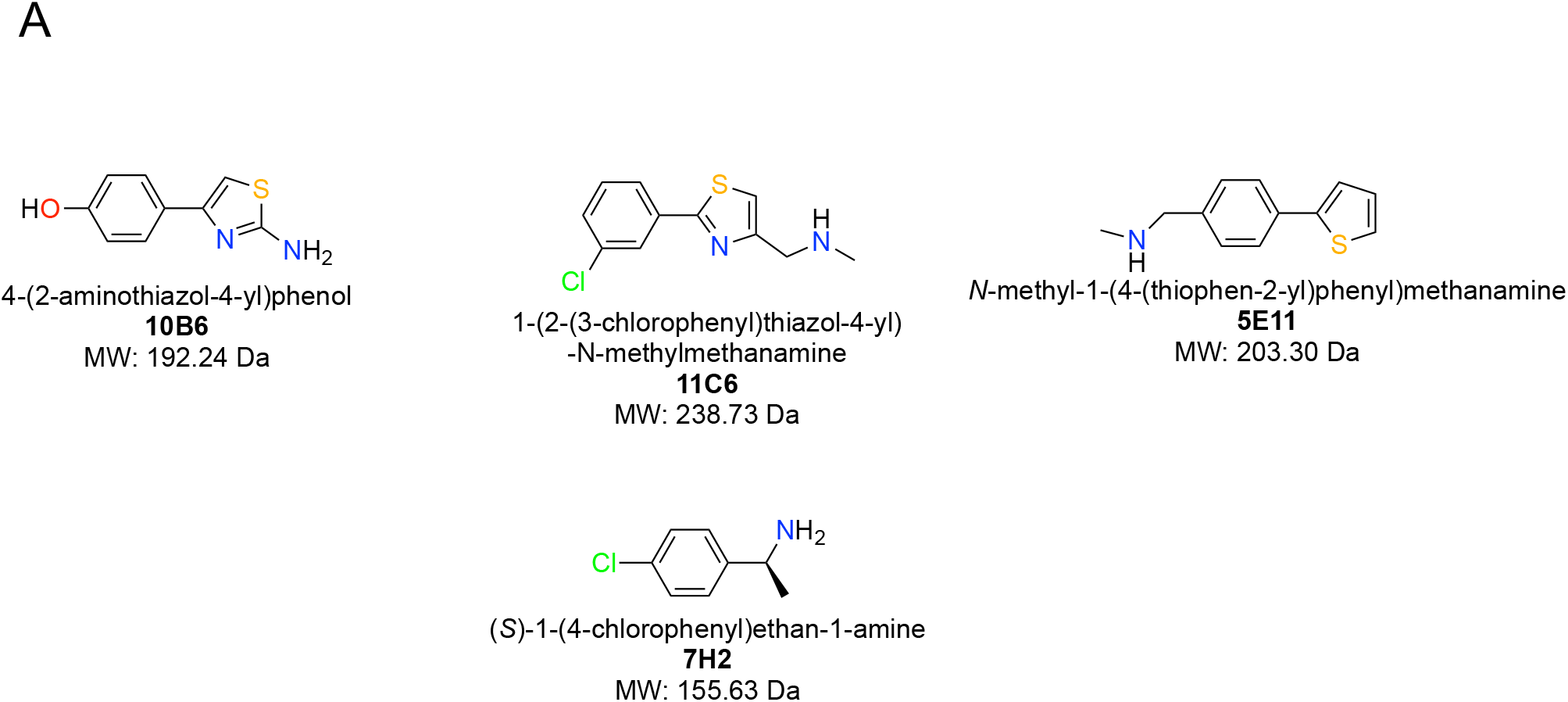
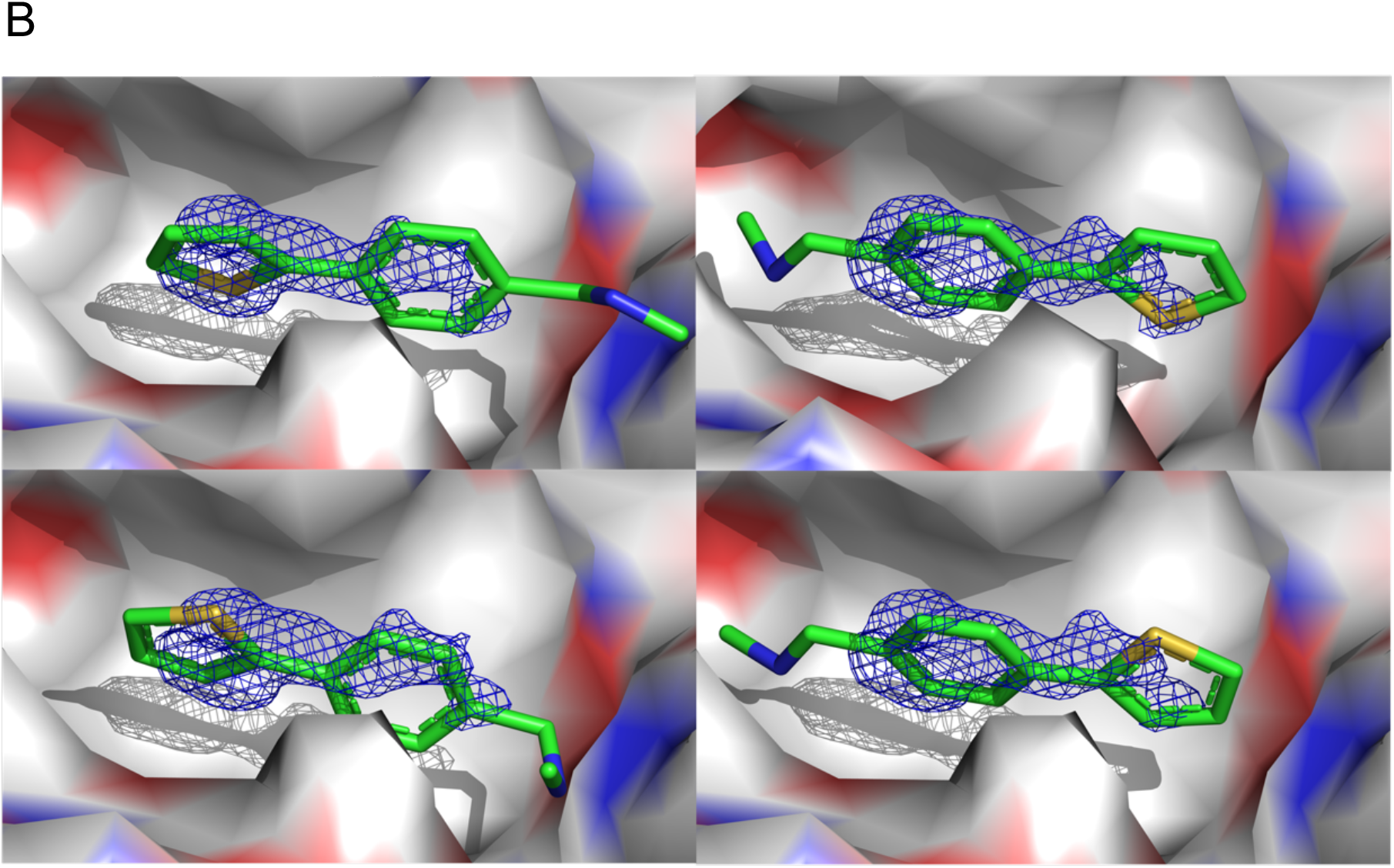
A) Chemical structures of the four nsp1-targeting fragment hits containing sulphur atoms and chloro substituents. B) Schematic representation of the four distinct orientations planar fragments can adopt in suboptimal real-life 2mF_o_-DF_c_ maps (rmsd = 1.0), exemplified by **5E11**. 180-degree rotations from one orientation to another can occur along horizontal or vertical axes, in particular when substituents are in quasi symmetric positions on one of the ring systems or if these possess some flexibility and are therefore not visible in the electron density.

To close this knowledge gap with respect to our project on SARS-CoV-2 nsp1, but also to establish a general procedure for similar cases, we used the long-wavelength beamline I23 at Diamond Light Source (DLS). Our aim was to clarify the exact orientation of four out of the six nsp1 fragment hits, containing sulphur and chlorine atoms by measuring diffraction data at wavelengths of 4.509 Å and 2.755 Å. The data collection at lower energy, just below the absorption edge of chloride, enables the detection of the exact location of anomalous scatterers i.e. sulphur and chloride, both present in our fragment hits, to be determined. I23 is a tuneable beamline where the X-ray diffraction experiment is optimised for data collection at very low energies (long wavelengths). At this beamline, the measurements are carried out in a vacuum environment using a multi-axis goniometer and it is equipped with a cylindrical shape detector ^11^.

Using sulphur and chlorine anomalous difference Fourier maps, we report the unambiguous identification of the binding orientation of four nsp1-targeting fragment hits. We show that one fragment is able to simultaneously bind in two distinct orientations albeit at distinct occupancies. Based on our results we propose a general procedure for unambiguously identifying binding orientations of 2-dimensional fragments containing either sulphur or chloro substituents as starting points for further optimisation for fragments, with substituents that can be examined within the wavelength range of most tuneable beamlines.

## Materials and Methods

### Expression, purification and crystallisation of SARS-CoV-2 nsp1_10-126_

The N-terminal domain of SARS-CoV-2 nsp1_10-126_, containing residues 10 to 126 and named nsp1 throughout the manuscript, was expressed, purified and crystallised as previously described^6^. The crystallisation condition used is 0.1 M BIS-TRIS pH 6.5, 0.2 M NaCl and 25% w/v Polyethylene glycol 3,350.

Fragment hits **5E11, 10B6**, and **11C6** were obtained from the Maybridge Ro3 library, while **7H2** was purchased from Molport (Cat. No.: HY-30379). Each of them was dissolved in DMSO at a concentration of 200 mM as a stock solution. 2 μL of each stock solution was mixed with 8 μL of the final buffer (10 mM HEPES pH 7.6 and 300 mM NaCl), giving a final concentration of 40 mM fragment containing 20% DMSO. Each fragment solution (1.5 μL) was added into approximately 1 μL of crystallisation drops, making a final fragment concentration of 24 mM and approximately 12% DMSO. These drops were incubated at room temperature for 4-5 h followed by crystal harvesting using loops, cryo-cooled in liquid nitrogen and stored in pucks for sample storage and shipment.

### Data collection, structure determination and refinement

The long-wavelength diffraction experiments were carried out at DLS, beamline I23. Each sample was subjected to X-rays at energies of 4.50 keV and 2.75 keV (corresponding to wavelengths of 2.755 Å and 4.509 Å), above and below the chlorine K absorption edge of 2.82 keV but both being above the sulphur K absorption edge of 2.47 keV. Diffraction experiments were performed in vacuum using the semi-cylindrical PILATUS 12M detector. During the measurements the temperature of protein crystals mounted on copper sample holders was estimated to be at ca. 80 K.

For each sample a 360-degree rotation and fine-slicing (0.10°) dataset was collected to obtain high multiplicity with the resolution only limited by detector dimensions (1.8 Å and 2.9 Å at wavelength 2.755 Å and 4.509 Å, respectively).

The data processing pipelines fast_DP and Xia2 [3.14.0], including Xia2_3dii and Xia2_dials, were automatically employed. PDB entry 8A55^12^ and the protein sequence were provided in ISPyB to trigger the processing pipeline Dimple [v2.6.2] to generate anomalous difference Fourier maps and MrBUMP to find a molecular replacement solution. Within the Dimple run, ten cycles of rigid-body refinement were performed, followed by four cycles of jelly body refinement and eight cycles of restrained refinement before starting to identify anomalous difference peaks^13^.

To identify unambiguous binding orientations of the four fragment hits, the nsp1-fragment coordinates and anomalous difference Fourier maps were overlaid with four nsp1 2mF_o_-DF_c_ and mF_o_-DF_c_ maps previously reported^6^ and inspected in COOT^14^. For **5E11, 10B6** and **11C6**, the binding site is located in proximity to residue Lys125, whereas for **7H2**, the shallower binding site is located adjacent to Pro109 as recently described^6^. Fragment hits were manually refitted into the ligand density with sulphur and (or) chlorine sitting in the centres of their anomalous difference Fourier map peaks. The updated structures were iteratively refined in Phenix^15^ and inspected in COOT. Fig 2 was prepared using Chemdraw^16^, whereas Fig 1 and Fig 3 to 6 were generated in Pymol^17^.

**Fig 2.**
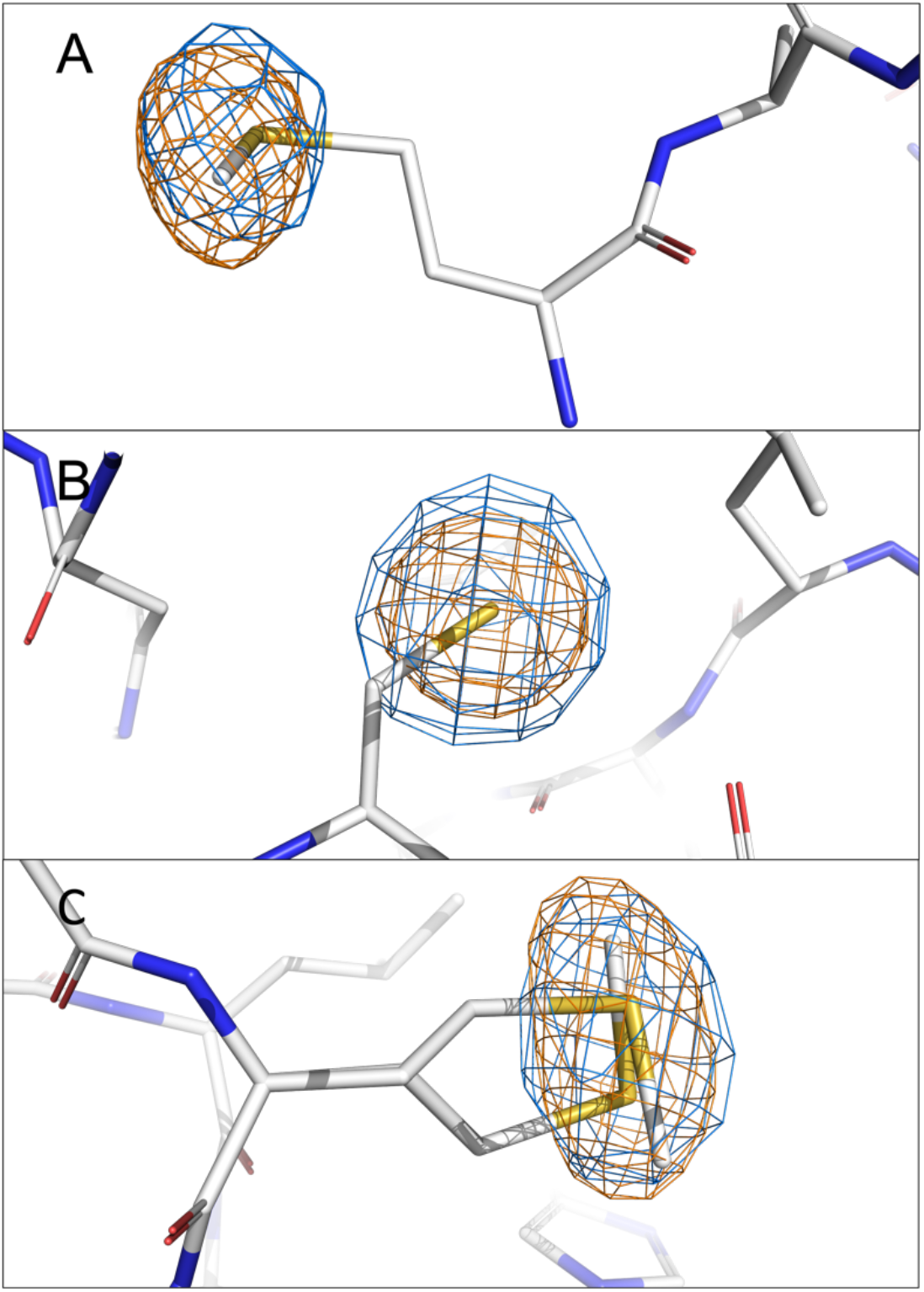
Validation of the localisation of anomalous signals of sulphur from A) Met9, B) Cys51 and C) Met85 in the anomalous difference Fourier maps (rmsd = 4.0) of nsp1-**7H2** complex. The anomalous peak at 4.50 keV is coloured in orange, while that at 2.75 keV is shown in marine. The two alternative conformations of Met85 are displayed and correlated well with the slightly elongated form of the anomalous signals.

**Fig 3.**
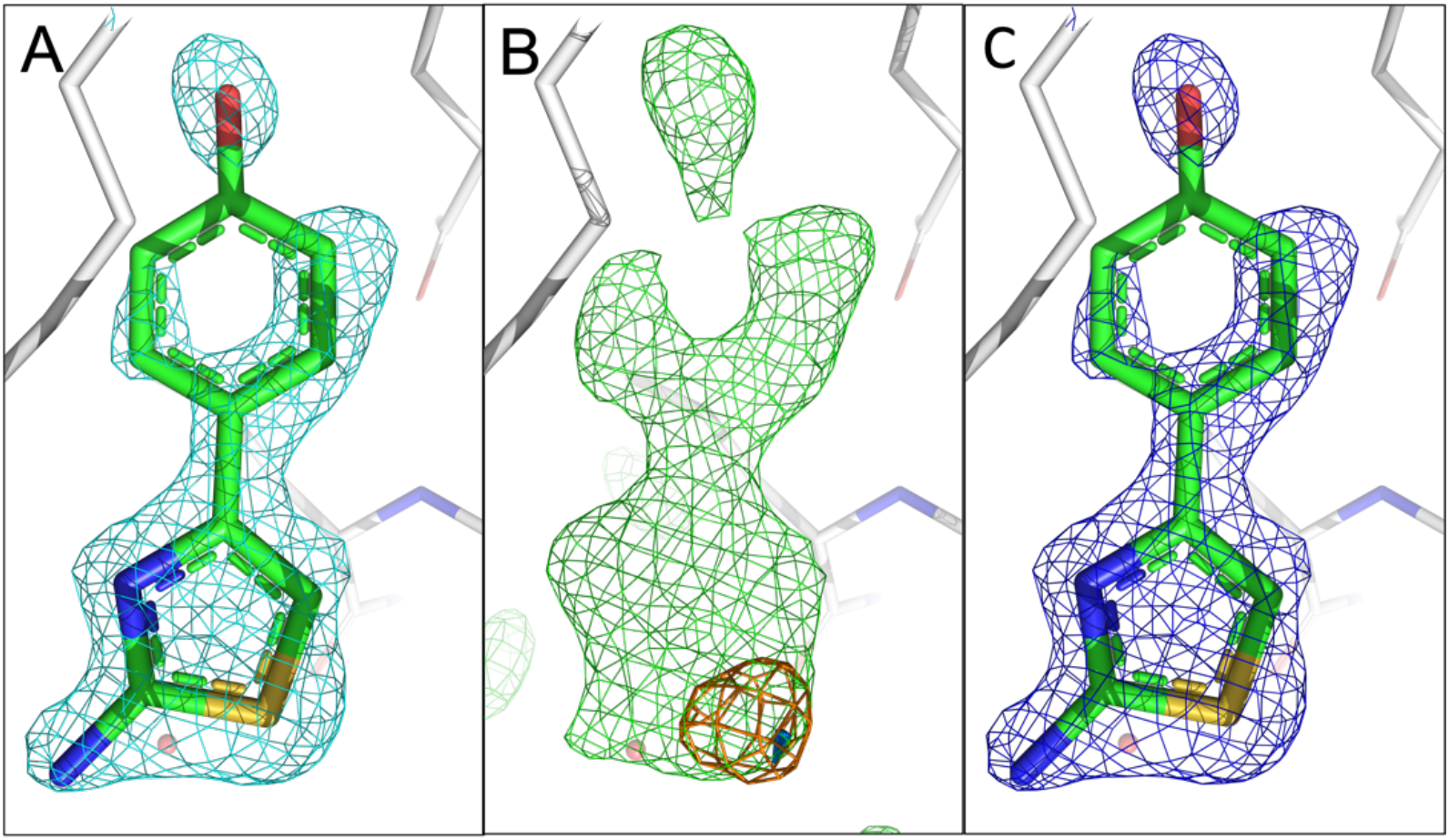
Comparison of the fragment binding site of previously reported conformation of **10B6** with those obtained using sulphur anomalous difference Fourier maps A) The published configuration of **10B6**. The 2mF_o_-DF_c_ map of the fragment is shown in cyan. **B)** Anomalous difference maps calculated from data collected at 4.5 keV (orange) and 2.75 keV (marine) with mF_o_-DF_c_ map (green) in the fragment region. **C)** Refined 2mF_o_-DF_c_ map of **10B6** with sulphur atom sitting in the centre of the anomalous peak. The rms deviations for the 2mF_o_-DF_c_, mF_o_-DF_c_ and anomalous difference Fourier maps are 1.0, 3.0, and 4.0, respectively, for figures 3 to 6. The electron density mostly accounts for **10B6**.

## Results

Data collection and refinement statistics are summarised in Tables 1 and 2. The very high I/σ*I* in the highest resolution shell (5.9 to 14.9) can in part be explained by the high resolution of the crystals but also with the low background during data collection. This is due to the complete absence of X-ray scattering in vacuum, which is typically a major cause of the background noise recorded at other macromolecular crystallography beamlines. First, we measured X-ray fluorescence spectra of the sample to identify chemical ions of interest that might be in the crystal or potentially be bound to the protein. The emission spectrum measured at excitation with 4.5 keV exemplified by the nsp1-**11C6** complex shows clear peaks assigned to K_a_ lines of sulphur and chlorine at 2.308 and 2.622 keV, respectively (Fig S1).

**Table 1.**
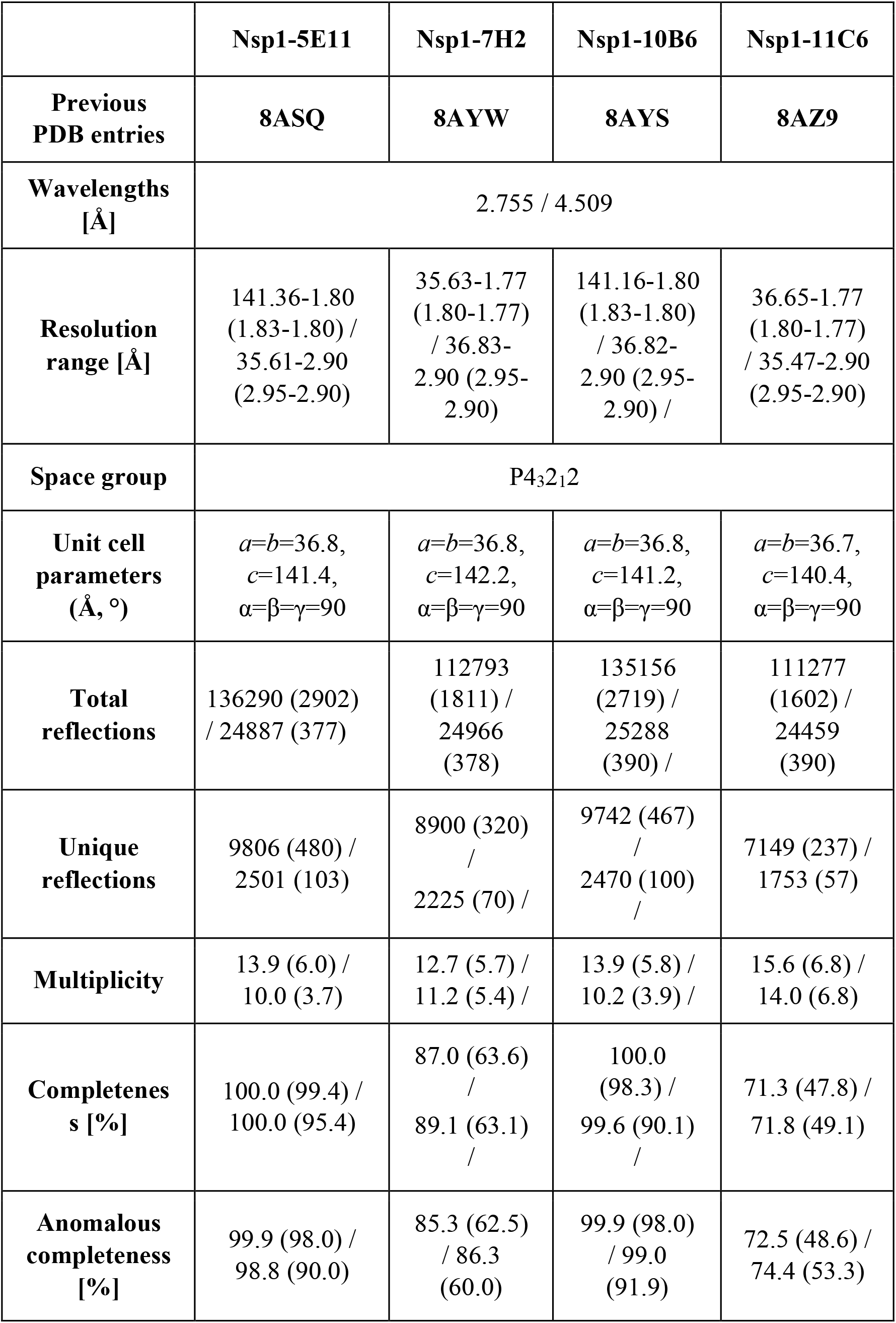
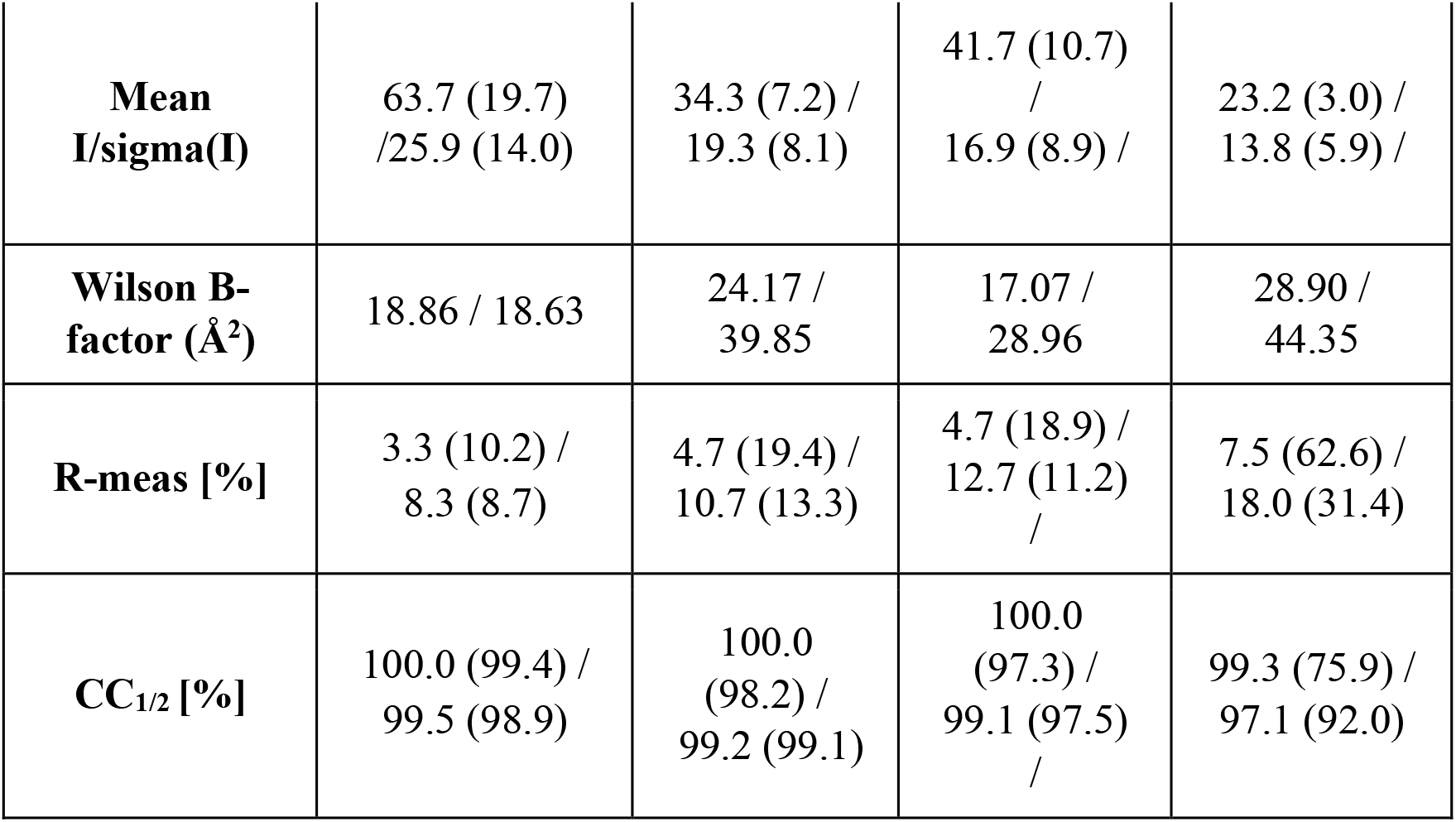
Data collection and refinement statistics for four nsp1-fragment hit complexes. measured at two distinct wavelengths. Data in the highest shell are shown in parenthesis.

|  | **Nsp1-5E11** | **Nsp1-7H2** | **Nsp1-10B6** | **Nsp1-11C6** |
| --- | --- | --- | --- | --- |
| **Previous PDB entries** | **8ASQ** | **8AYW** | **8AYS** | **8AZ9** |
| **Wavelengths [Å]** | 2.755 / 4.509 |  |  |  |
| **Resolution range [Å]** | 141.36-1.80<br>(1.83-1.80) /<br>35.61-2.90<br>(2.95-2.90) | 35.63-1.77<br>(1.80-1.77)<br>/ 36.83-2.90 (2.95-2.90) | 141.16-1.80<br>(1.83-1.80)<br>/ 36.82-2.90 (2.95-2.90) / | 36.65-1.77<br>(1.80-1.77)<br>/ 35.47-2.90<br>(2.95-2.90) |
| **Space group** | P4<sub>3</sub>2<sub>1</sub>2 |  |  |  |
| **Unit cell parameters (Å, °)** | $a=b=36.8$,<br>$c=141.4$,<br>$\alpha=\beta=\gamma=90$ | $a=b=36.8$,<br>$c=142.2$,<br>$\alpha=\beta=\gamma=90$ | $a=b=36.8$,<br>$c=141.2$,<br>$\alpha=\beta=\gamma=90$ | $a=b=36.7$,<br>$c=140.4$,<br>$\alpha=\beta=\gamma=90$ |
| **Total reflections** | 136290 (2902)<br>/ 24887 (377) | 112793<br>(1811) /<br>24966<br>(378) | 135156<br>(2719) /<br>25288<br>(390) / | 111277<br>(1602) /<br>24459<br>(390) |
| **Unique reflections** | 9806 (480) /<br>2501 (103) | 8900 (320)<br>/<br>2225 (70) / | 9742 (467)<br>/<br>2470 (100)<br>/<br>/ | 7149 (237) /<br>1753 (57) |
| **Multiplicity** | 13.9 (6.0) /<br>10.0 (3.7) | 12.7 (5.7) /<br>11.2 (5.4) / | 13.9 (5.8) /<br>10.2 (3.9) / | 15.6 (6.8) /<br>14.0 (6.8) |
| **Completeness [%]** | 100.0 (99.4) /<br>100.0 (95.4) | 87.0 (63.6)<br>/<br>89.1 (63.1)<br>/<br>/ | 100.0<br>(98.3) /<br>99.6 (90.1)<br>/<br>/ | 71.3 (47.8) /<br>71.8 (49.1) |
| **Anomalous completeness [%]** | 99.9 (98.0) /<br>98.8 (90.0) | 85.3 (62.5)<br>/ 86.3<br>(60.0) | 99.9 (98.0)<br>/ 99.0<br>(91.9) | 72.5 (48.6) /<br>74.4 (53.3) |
| **Mean I/sigma(I)** | 63.7 (19.7) / 25.9 (14.0) | 34.3 (7.2) / 19.3 (8.1) | 41.7 (10.7) / 16.9 (8.9) / | 23.2 (3.0) / 13.8 (5.9) / |
| **Wilson B-factor (Å<sup>2</sup>)** | 18.86 / 18.63 | 24.17 / 39.85 | 17.07 / 28.96 | 28.90 / 44.35 |
| **R-meas [%]** | 3.3 (10.2) / 8.3 (8.7) | 4.7 (19.4) / 10.7 (13.3) | 4.7 (18.9) / 12.7 (11.2) / | 7.5 (62.6) / 18.0 (31.4) |
| **CC<sub>1/2</sub> [%]** | 100.0 (99.4) / 99.5 (98.9) | 100.0 (98.2) / 99.2 (99.1) | 100.0 (97.3) / 99.1 (97.5) / | 99.3 (75.9) / 97.1 (92.0) |

**Table 2.** Refinement statistics for three nsp1-fragment complexes. Mtz files used to generate updated nsp1-fragment complexes are from previous data collections. Data in the highest shell are shown in parenthesis.

|  | **Nsp1-5E11** | **Nsp1-7H2** | **Nsp1-11C6** |
| --- | --- | --- | --- |
| **PDB ID** | **8CRF** | **8CRK** | **8CRM** |
| **R<sub>cryst</sub>/R<sub>free</sub> [%]** | 19.2 (30.8) / 24.4 (34.6) | 16.8 (26.8) / 19.2 (27.7) | 17.9 (29.2) / 20.4 (28.9) |
| **Reflections used in refinement** | 34387 (3461) | 40247 (3890) | 19211 (1871) |
| **Reflections used for R-free** | 868 (87) | 988 (96) | 951 (97) |
| **Total no. of non-hydrogen atoms (protein)** | 1013 | 1041 | 1113 |
| **No. of protein/ligand/solvent atoms** | 885 / 54 / 100 | 919 / 20 / 112 | 973 / 26 / 125 |
| **Average B-factor/protein/ligands/solvent** | 26.4 / 23.8 / 75.8 / 35.6 | 22.6 / 21.1 / 48.1 / 32.6 | 25.3 / 23.9 / 54.9 / 32.9 |
| **RMSD (bonds, angles)** | 0.012 / 1.30 | 0.012 / 1.39 | 0.012 / 1.27 |
| **Ramachandran favoured/allowed/outliers/rotamer outliers [%]** | 99.1 / 0.90 / 0 / 1.1 | 97.4 / 2.7 / 0 / 1.00 | 96.6 / 3.5 / 0 / 0.9 |
| **Clashscore** | 5.5 | 5.3 | 6.0 |

To validate our system of locating the chlorine and sulphur atoms in anomalous difference Fourier maps at two distinct energies, the anomalous difference Fourier maps of sulphur from methionine and cysteine side chains present in nsp1 were visually inspected. There are two methionines and one cysteine in the N-terminal domain of nsp1. The anomalous signals originating from the sulphur atoms in these residues are visible in both 4.5 keV and 2.75 keV anomalous difference Fourier maps except for nsp1-**11C6**, for which the anomalous signal of sulphur in Met9 was not detected at 2.75 keV. This is possibly because of the low completeness of the specific data set and the flexibility of Met9 as the first residue at the N-terminus. The signal levels of the sulphur atoms in these residues for all four datasets are listed in Table 3. The overlapping local electron density maps for these residues in the nsp1-**7H2** complex at two diffraction energies (Fig 2 have been selected as an example. Interestingly, the first amino acid Met9 was previously placed in a somewhat uncertain position due to its side chain flexibility and therefore lack of electron density for certain side chain atoms. However, the anomalous signal of the sulphur atom clearly reveals the correct localisation of the sulphur atom (Fig 2A). There is only a single position for the Cys51 side chain, supported by the spherical electron density of the anomalous signal (Fig 2B). The two alternative conformations of the Met85 side chain built in the previous complex (PDB entry 8AYW) structure are also clearly supported by the elongated anomalous sulphur signal (Fig 2C). These results not only validate the reliability of our approach but also show the significance of anomalous difference Fourier maps in revealing missing details in native (non-anomalous) electron-density maps.

**Table 3.** Anomalous signal levels of sulphur from Met9, Cys51 and Met85 in four nsp1-fragment datasets measured at two distinct diffraction energies.

| Fragment hits |  | 5E11 | 7H2 | 10B6 | 11C6 |
| --- | --- | --- | --- | --- | --- |
| Diffraction energy (keV) |  | 2.75 / 4.50 |  |  |  |
| **S ($\sigma$)** | **Met9** | 8.0 / 12.7 | 6.1 / 11.1 | 8.0 / 14.2 | n.d. / 7.4 |
|  | **Cys51** | 18.8 / 46.8 | 12.7 / 34.0 | 14.02 / 38.6 | 7.3 / 21.5 |
|  | **Met85** | 10.6 / 21.6 | 7.4 / 17.3 | 8.5 / 18.4 | 5.3 / 13.1 |

\*n.d. not detected

Fragment **10B6** (Fig 1A) consists of two rings, containing a heterocyclic 5- and an aromatic 6-ring system. It contains a single sulphur atom in the 2-aminothiazol ring. Only a single anomalous peak is visible at the binding pocket in the 4.5 KeV anomalous difference Fourier maps at a contour level of 4.0 rmsd. Having inspected the measured anomalous difference Fourier map we concluded that a value of > 4.0 can be adopted as a positive indication of the presence of the atom. The signal is still visible in the 2.75 KeV anomalous difference Fourier maps (rmsd of 4.0), albeit a smaller peak, confirming the existence of sulphur in fragment **10B6**, corresponding to its chemical structure. We could easily fit **10B6** into the density with the sulphur atom sitting in the centre of the sulphur peak. A comparison with our recently published fragment configuration shows that our previous assumption was correct, and we only observed a single configuration in the difference map (Fig 3 A-C).

Fragment **11C6** can be considered as a closely related analogue of **10B6** (Fig 1A). The sulphur atom is located in a thiazole ring system, and it also contains a chloro substituent in the *meta*-position of the phenyl group, allowing us to observe two anomalous signals in the experimental anomalous difference Fourier map. Indeed, two anomalous peaks are visible in the fragment binding site in the 4.5 KeV anomalous difference Fourier maps (rmsd of 4.0). Neither of the signals is visible in the 2.75 KeV anomalous difference Fourier maps even at contour level of 3.0 rmsd, which is expected for the chlorine-but not for the sulphur atom in **11C6**. This could be caused by the lower occupancy of **11C6** in the binding site or the overall weak anomalous signals at lower energy maps as observed for sulphur anomalous signals from Met9. However, as one anomalous peak is located approximately at an extension of the ligand density while the second one is slightly away from the main ligand density, we can easily distinguish and allocate the sulphur-containing ring as well as the chloro substituent, making unambiguous fragment orientation and conformation possible (Fig 4 A-C).

**Fig 4.**
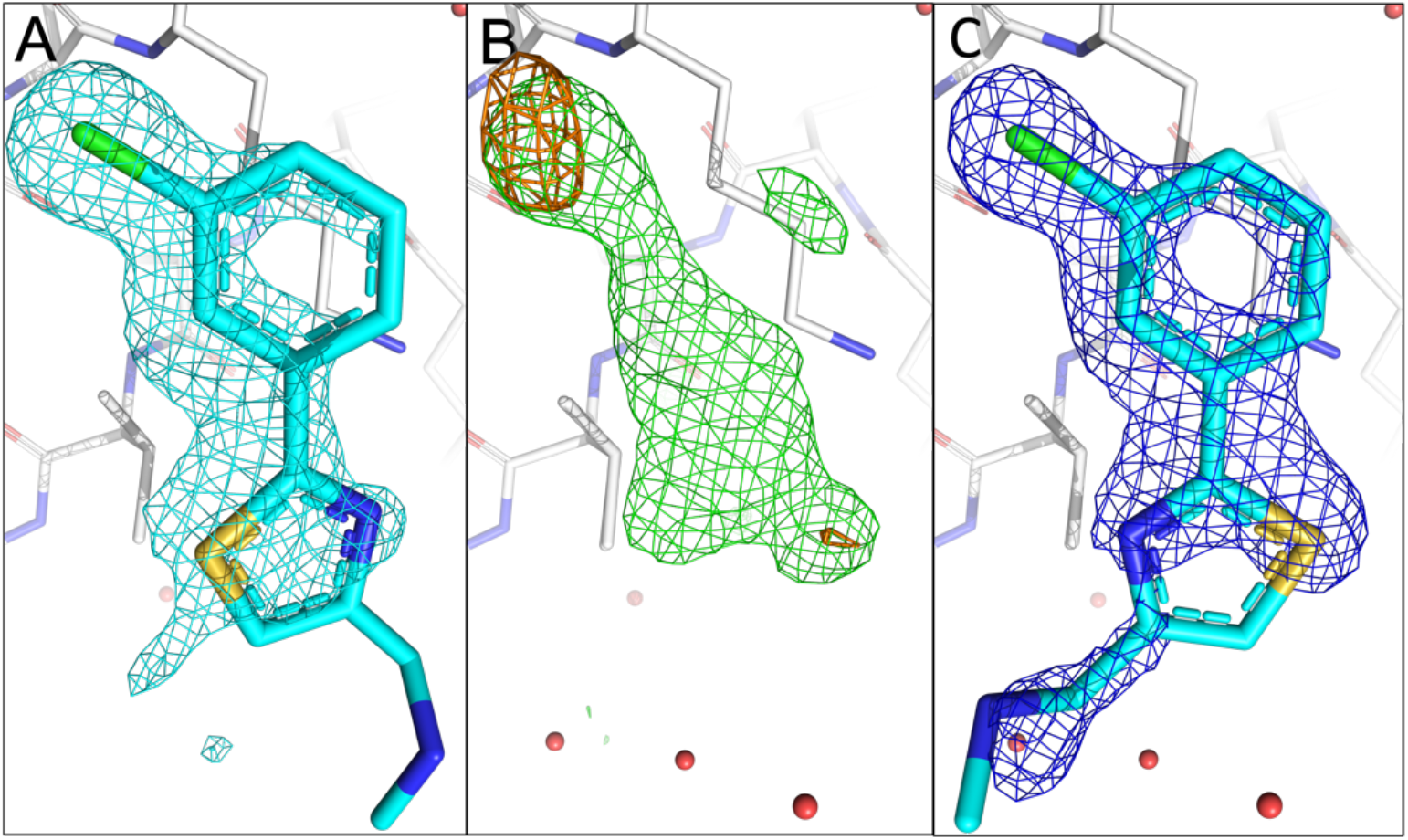
Comparison of the fragment binding site of the previously reported conformation of **11C6** with that obtained using sulphur and chlorine anomalous difference Fourier maps. **A)** The published configuration of **11C6**. The 2mF_o_-DF_c_ map of the fragment is shown in cyan. **B)** Anomalous maps calculated from data collected at 4.5 keV (orange) with mF_o_-DF_c_ map (green) in the fragment binding region. The anomalous peak from 2.75 keV data does not appear in the sulphur location possibly because of the low occupancy of **11C6. C)** Refined 2mF_o_-DF_c_ map of **11C6** with the chlorine and sulphur atoms sitting in the centres of the anomalous peaks. The electron density partially covers **11C6**, but the orientation of the fragment hit could be confirmed by the anomalous signals for sulphur and chlorine.

Compared to the previous two fragments hits, **5E11** (Fig 1A) represents a much simpler fragment structure containing a single N-methyl-methanamine substituent in the *para*-position of the phenyl ring system with the sulphur atom being part of a thiophen ring system. To our surprise, for **5E11**, two anomalous peaks are visible at the fragment binding site in the 4.5 KeV anomalous difference Fourier maps (rmsd of 4.0), albeit with different intensities. The two signals are still visible in the 2.75 KeV anomalous difference Fourier maps (rmsd of 3.0), confirming the existence of sulphur in **5E11 (Fig S2)**, in agreement with its chemical structure. However, this also implies two simultaneous binding conformations with distinct occupancies. In our previous work, we had fitted **5E11** in a single orientation as the electron density even at very high resolution did not reveal multiple orientations. However, our new more detailed approach employed here clearly reveals two orientations for the sulphur atom. Therefore, we manually fitted **5E11** in two distinct orientations with different occupancy (Fig 5 C). This is an excellent example of how to use anomalous difference data to identify all possible binding orientations of a fragment before engaging in SAR-by-catalogue or by synthesising novel fragment analogues.

**Fig 5.**
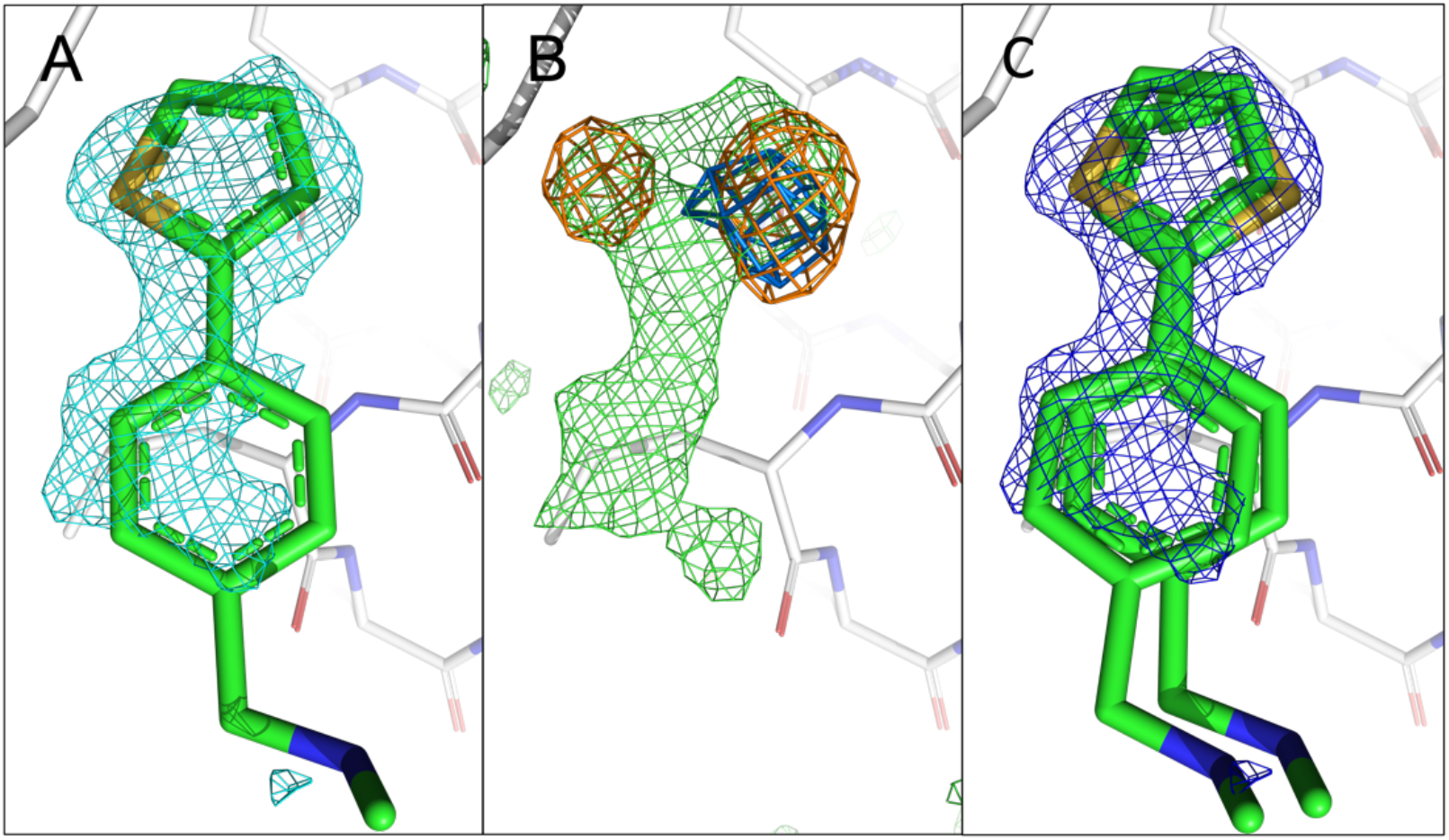
Comparison of the previously reported single conformation of **5E11** with those obtained using sulphur anomalous difference Fourier maps. **A)** The published configuration of **5E11**. The 2mF_o_-DF_c_ map of the fragment is shown in cyan. **B)** Anomalous difference Fourier maps calculated from datasets collected at 4.5 keV (orange) and 2.75 keV (marine) with mF_o_-DF_c_ map (green) in the fragment region. **C)** Refined 2mF_o_-DF_c_ map of **5E11** in the two orientations with sulphur sitting in the centres of the two anomalous peaks obtained from 4.5 keV data. The electron density mostly covers **5E11** in both distinct orientations, but there is no density around the N-methyl-methanamine substituent, indicating its flexibility and complicating unambiguous placement of the fragment in the absence of sulphur peaks.

Whereas the previous three fragment hits can be considered as chemically closely related analogues, **7H2** represents a significantly smaller fragment of 155 Da containing a single central phenyl ring with two distinct substituents binding to a shallower site on nsp1 (Fig 6A-C). Due to its quasi-symmetrical chemical structure (Fig 1A), a rotation around its long axis does not lead to a new distinct orientation, as the ethan-1-amine substituent contains a rotatable bond. However, a vertical axis through its phenyl ring will lead to two distinct orientations with each of the two substituents changing positions. For **7H2**, only one anomalous peak at the binding pocket is visible in the 4.5 KeV anomalous difference Fourier maps at rmsd of 4.0. The peak disappears in the 2.75 KeV anomalous difference Fourier maps (same rmsd level), consistent with the existence of a single chlorine in **7H2**. Consequently, we fitted **7H2** into the density with the chloro substituent located at the centre of the anomalous peak, in the opposite direction of our previously published orientation.

**Fig 6.**
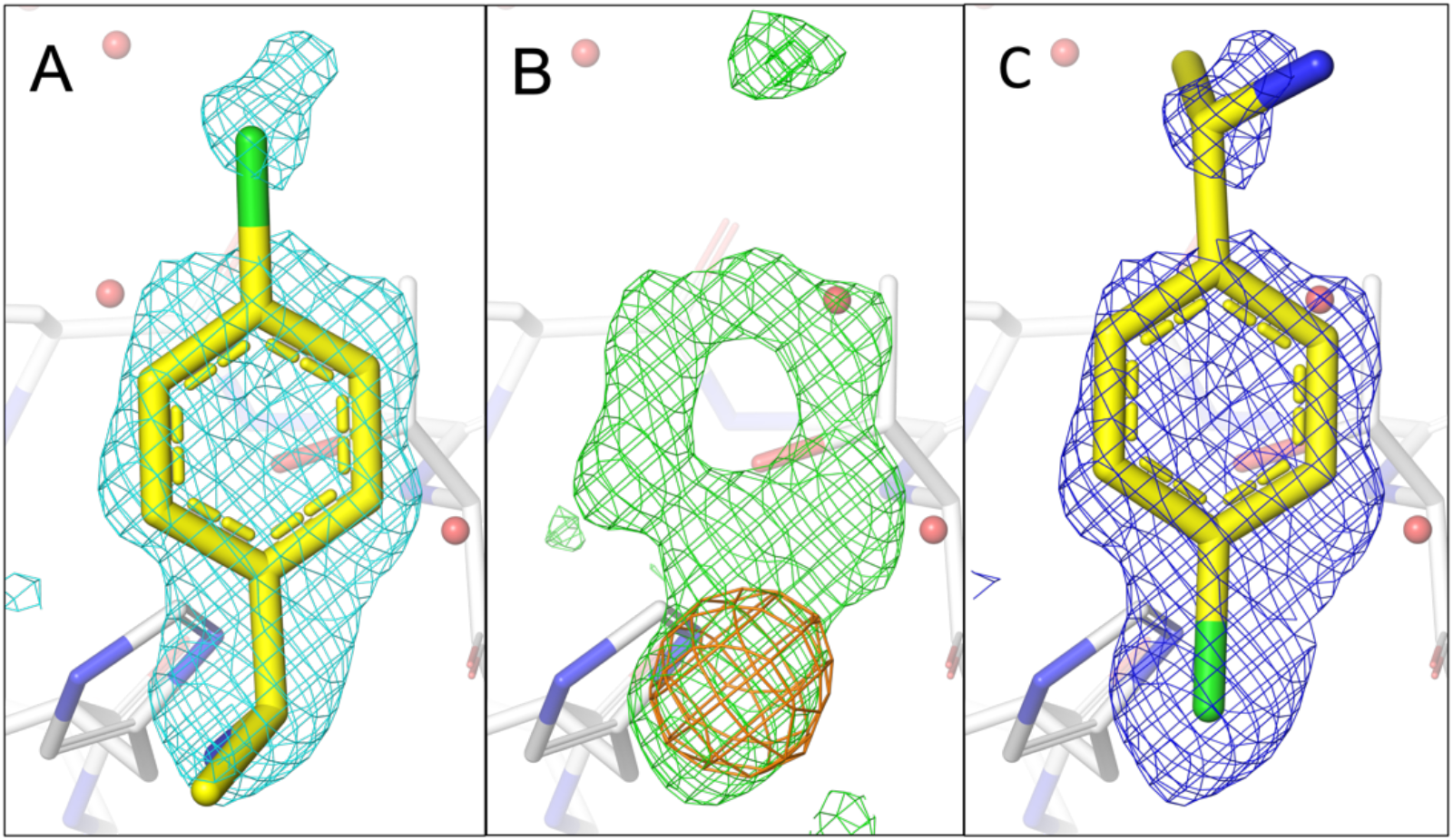
Comparison of the fragment binding site of previously reported conformation of **7H2** with that obtained using a chlorine anomalous difference Fourier map. **A)** The published configuration of **7H2** with the 2mF_o_-DF_c_ map around the fragment is shown in cyan. **B)** Anomalous difference Fourier maps calculated from data collected at 4.5 keV (orange) with the mF_o_-DF_c_ map (green) in the fragment binding region. **C)** Refined 2mF_o_-DF_c_ of **7H2** with the chlorine located in the centre of the anomalous peak. The electron density mostly covers **7H2**.

## Discussion and Conclusions

The unambiguous identification of atomic elements from phased anomalous Fourier difference maps relies on the measurements above and below an absorption edge of the atom. This technique is shown to be very useful in unravelling the structure and function of ion-binding proteins, but it has been challenging for the lighter atomic weight elements at standard synchrotron beamlines due to air absorption as well as other technical constraints^18^. The DLS I23 beamline operates in vacuum and is optimised for the data collection at wavelengths up to 5.9 Å enabling it to reach K-absorption edges of P, S, Cl, K, and Ca (see Fig. S1). The absence of scattering from the air results in a very low endogenous background of the detector that translates into the improved signal-to-noise ratio in the anomalous difference Fourier maps. Furthermore, due to a much enhanced value of f’’ at L-absorption edges of other biologically relevant halides, i.e. Br and I substituents, the identification of their position is more straightforward when measured at I23^19^. Consequently, by conducting experiments around the absorption edges of interest located in this energy range one can determine the identity and location of relevant substituents in fragments that bind to proteins and then use this information for fitting in the electron density map.

A previous study revealed that less than 2% of entries containing small molecules/ligands reported two or more alternate conformations^1^, which is probably a consequence of past limitations such as beamline design, a vacuum environment to reduce noise and dedicated detectors. The available published information on ligand orientation or conformation is even rarer for fragment hits with molecular weights of < 300 Da. The extent of simultaneously existing multiple orientations of fragments binding to the same site may be even more frequent as their molecular weight is significantly smaller than that of small molecules with suboptimal interactions with residues in the binding pocket at the start of a campaign. Planar fragments that lack 3D characteristics, or those with low occupancy and/or flexible sub-optimised ligands can now be confidently placed. The “flatness” of fragments has been recognised as a challenge and various attempts have been made to incorporate fragments with 3D characteristics into libraries^20, 21^. Therefore, it is not surprising that studies involving anomalous difference maps for fragments are rare compared to small molecule work. Brominated fragments have been preferentially employed, likely due to their easy use as they can be measured using even x-ray generators and do not require tuneable beamlines. For example, in a pilot study, brominated fragments have been employed to identify fragment hits at low resolution ^22^. A library with 68 brominated fragments was screened against HIV protease to successfully identify binding sites and fragment orientations ^23^. In another project the authors employed brominated and chlorinated fragment analogues using a tuneable beamline to locate novel fragment binding sites in glycerol-3-phosphate dehydrogenase and to identify two distinct binding orientations^24^. In our work, we provide an optimised protocol using a dedicated tuneable beamline to focus on sulphur and chlorine containing substituents in fragment hits. The long-wavelength MX beamline I23 operating in a vacuum environment has made this type of study possible, allowing for experimentation in proximity to the absorption edges of sulphur and chlorine. Additionally, this controlled setting effectively reduces noise levels, optimizing the detection of anomalous difference signals.

A recent study showed that 24% of marketed drugs contained sulphur atoms and approximately 28% were halogenated ^25^. Halogen-enriched fragment libraries with fragments containing at least one halogen atom per fragment are available^26^. We probed recently identified fragment hits obtained by fragment-based screening via X-ray crystallography^6^. The resulting fragment hits were predominantly planar due to aromatic ring systems and despite working at resolutions between 1.1 and 1.4 Å, electron density maps remained ambiguous, even after extensive refinement. Not unexpectedly, neither R_free_ values, density maps or atomic B-factors were fully conclusive raising our suspicion of the presence of multiple fragment orientations. Intuition led to the correct placement of two out of four fragment hits, but intuition should not be the basis for subsequent rational structure-based design. Consequently, we identified fragment **5E11** that simultaneously binds to nsp1 in two distinct orientations as revealed by two peaks in the anomalous difference Fourier maps. Based on their intensity we could also estimate the occupancy for each orientation with a preference of the sulphur atom in the thiophene ring pointing towards the protein surface and establishing hydrogen bond interaction with Lys47^6^ rather than pointing towards the solvent. As both sulphur difference peaks are located on the same ring, our approach is powerful enough to pinpoint individual atoms on a 5-membered ring system with a distance of only 2.58 Å between them. A further improvement of our approach could be to measure a third dataset at a shorter wavelength to further improve data resolution.

Based on our work we recommend the following procedure for data collection of planar fragments with ambiguous or weak electron density due to low occupancy or the presence of multiple orientations. First collect data at the highest possible resolution. If doubts exist about the correct orientation or conformation of the fitted fragment making rational fragment optimisation a challenge, use fragment analogues with detectable substituents such as sulphur and chlorine by measuring anomalous difference data above and below the absorption edge of the atom at a tuneable beamline to clearly locate it orientation. If radiation damage is not an issue, data collection can include a native dataset at high resolution measured on the same crystal. Although currently limited to sulphur and chlorine-containing fragments, work is currently in progress to expand our approach to other elements frequently present in fragments.

## Abbreviations

BIS-TRIS: Bis(2-hydroxyethyl)amino-tris(hydroxymethyl)methane
CoV: Coronavirus
Covid-19: Coronavirus Disease 2019
Da: Dalton
DMSO: Dimethyl sulfoxide
DLS: Diamond Light Source
FBDD: Fragment-based drug discovery
HEPES: (4-(2-hydroxyethyl)-1-piperazineethanesulfonic acid
IPTG: Isopropyl-β-D-thiogalactopyranoside
mRNA: Messenger ribonucleic acid
Nsp: Non-structural protein
PDB: Protein data bank
PMSF: Phenylmethylsulfonyl Fluoride
rmsd: Root Mean Square Deviation
SARS-CoV-2: Severe Acute Respiratory Syndrome Coronavirus

## Supplementary materials

**Fig. S1.**
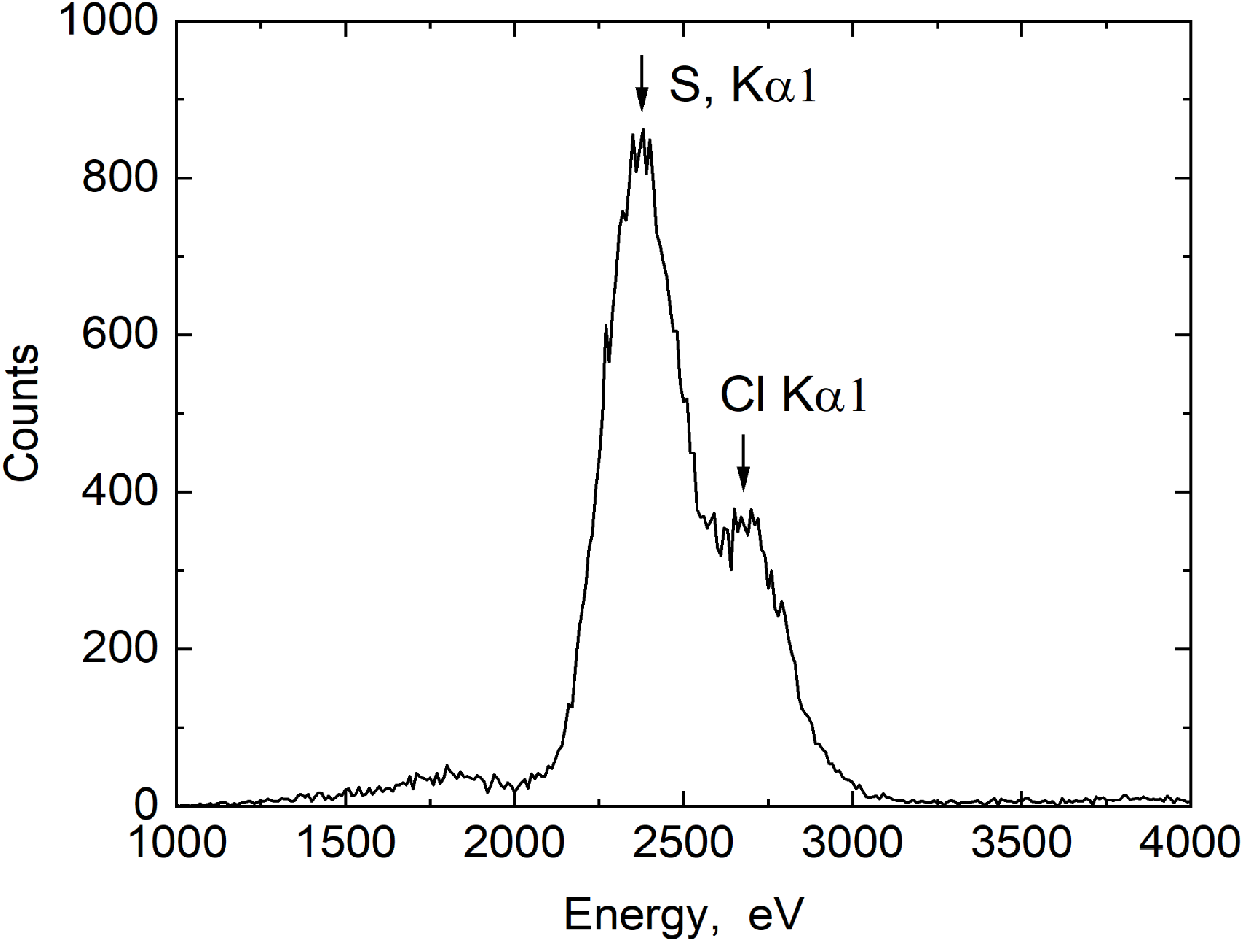
X-ray fluorescence spectra of the nsp1-**11C6** crystal sample. The peaks at 2308 and 2622 keV correspond to K_a1_ emission lines of sulphur and chlorine. The elevation at low energy is a background predominantly caused by X-ray emission of silicon from the detector.

**Fig. S2.**
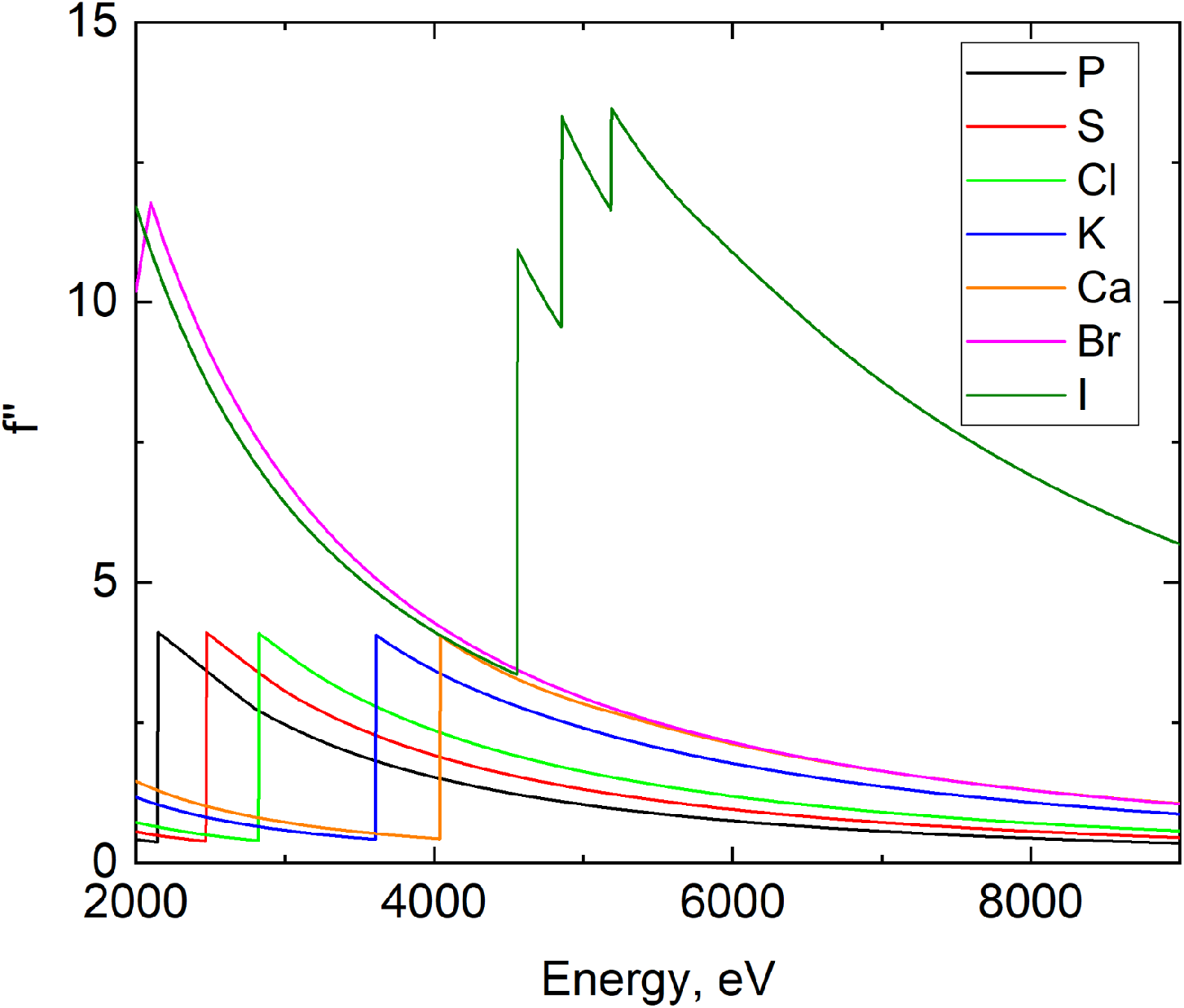
Energy dependence of the anomalous contribution to the structural factor f’’ for biologically relevant light ions and halides (data from www.bmsc.washington.edu). The value of f’’ manifests a sharp increase at the K-edge of light atoms (P, S, Cl, K, Ca) or L-edges of Br and I thus enabling positive identification of the ions.

## Acknowledgements

This publication contains part of the doctoral thesis of Shumeng Ma.

